# Omicron and Delta Variant of SARS-CoV-2: A Comparative Computational Study of Spike protein

**DOI:** 10.1101/2021.12.02.470946

**Authors:** Suresh Kumar, Thiviya S. Thambiraja, Kalimuthu Karuppanan, Gunasekaran Subramaniam

## Abstract

Emerging SARS-CoV-2 variants, especially those of concern, may have an impact on the virus’s transmissibility and pathogenicity, as well as diagnostic equipment performance and vaccine effectiveness. Even though the SARS-CoV-2 Delta variant (B.1.617.2) emerged during India’s second wave of infections, Delta variants have grown dominant internationally and are still evolving. On November 26, 2021, WHO identified the variant B.1.1.529 as a variant of concern, naming it Omicron, based on evidence that Omicron contains numerous mutations that may influence its behaviour. However, the mode of transmission and severity of the Omicron variant remains unknown. We used computational studies to examine the Delta and Omicron variants in this work and found that the Omicron variant had a higher affinity for human ACE2 than the Delta variant due to a significant number of mutations in the SARS-CoV-2 receptor binding domain, indicating a higher potential for transmission. Based on docking studies, the Q493R, N501Y, S371L, S373P, S375F, Q498R, and T478K mutations contribute significantly to high binding affinity with human ACE2. In comparison to the Delta variant, both the entire spike protein and the RBD in Omicron include a high proportion of hydrophobic amino acids such as leucine and phenylalanine. These amino acids are located within the protein’s core and are required for structural stability. Omicron has a higher percentage of alpha-helix structure than the Delta variant in both whole spike protein and RBD, indicating that it has a more stable structure. We observed a disorder-order transition in the Omicron variant between spike protein RBD regions 468-473, and it may be significant in the influence of disordered residues/regions on spike protein stability and binding to ACE2. A future study might investigate the epidemiological and biological consequences of the Omicron variant.

## Introduction

SARS-CoV-2 (Severe acute respiratory syndrome coronavirus type 2) is a coronavirus that caused the Covid-19 disease outbreak in late 2019 in Wuhan China. By early 2020, the disease had rapidly spread across the world and was declared a global pandemic as a public health emergency of international concern. The virus spreads from person to person by respiratory droplets in close contact between sick and asymptomatic people (within 6 feet) ^1^. Transmission by aerosols and maybe contact with fomites is also a possibility, although this is not considered to be the most probable route ^2^. SARS-CoV-2 pathogenesis is dependent on the viral spike protein binding to angiotensin-converting enzyme 2 (ACE2) receptors, with cell entrance required ACE2 receptor cleavage by a type 2 transmembrane serine protease to activate the viral spike protein ^3^. COVID-19 individuals have a wide range of clinical symptoms, from moderate to severe, fast progressive, and acute disease ^4^. The diagnosis of COVID-19 is non-specific, and the virus may manifest itself in a variety of ways, ranging from no symptoms (asymptomatic) to severe pneumonia and death. The CoVID-19 pandemic response plan is based on the development of therapeutic alternatives and vaccination formulations ^5-7^.

The term “variant of concern” (VOC) for SARS-CoV-2 (which produces COVID-19) refers to viral variants with mutations in their spike protein receptor-binding domain (RBD) that dramatically improve binding affinity in the RBD-hACE2 complex while also causing fast dissemination in human populations ^8^. Increased viral replication increases the likelihood of SARS-CoV-2 mutations forming. Therefore, the only option to end the pandemic is for effective vaccinations against circulating variations to be extensively and fairly delivered globally. Because raising nations are rushing to vaccinate their people within months, they risk SARS-CoV-2 evolving into a new lineage that vaccines may not be able to protect against in other countries. To combat some emerging SARS-CoV-2 strains, new vaccinations may need to be developed regularly. With the introduction of extremely infectious SARS-CoV-2 variants, greater vaccine penetration will be required to build protective immunity, and children may also need to be vaccinated ^9^.

While the majority of SARS-CoV-2 sequence changes are projected to be detrimental and swiftly removed or to be neutral, a small number are predicted to influence functional characteristics, possibly modifying infection rate, disease severity, or interactions with the host immune system ^10^. Nonetheless, beginning in late 2020, the development of SARS-CoV-2 has been marked by the introduction of ‘variants of concern,’ or changes in viral properties such as disease transmission and antigenicity, most likely because of the changing immunological composition of the human species.

The delta variant (B.1.617.2) was discovered for the first time in India in late 2020. The Delta version may have invaded over 163 nations by August 24, 2021. The World Health Organization (WHO) stated in June 2021 that the Delta strain is on its way to becoming the most prevalent strain in the world ^11^. Therefore, the Delta variant was changed from Variant of Interest (VOI) to Variant of Concern (VOC). According to present evidence, the SARS-CoV-2 Delta variant of concern (VOC) is 40-60% more transmissible than the Alpha (B.1.1.7) VOC and may be associated with an increased risk of hospitalisation. The Delta VOC mostly endangers those who are unvaccinated or just partially vaccinated ^12^.

On November 26, 2021, the World Health Organization’s Technical Advisory Group on Virus Evolution (TAG-VE) proposed that variant B.1.1.529, commonly known as Omicron, be identified as a variant of concern. The TAG-VE made this decision after discovering that Omicron has several mutations that might impact how quickly it spreads or the severity of the disease it causes. The spike protein’s variation is determined by thirty mutations, 15 of which occur in the receptor-binding domain, as well as three small deletions and one minor insertion. This mutation was discovered in samples collected in Botswana on November 11, 2021, and South Africa on November 14, 2021. As of November 26, 2021, travel-related occurrences have also been documented in Belgium, Hong Kong, and Israel. The Omicron variant is the most divergent strain seen in significant numbers so far during the pandemic, raising concerns that it may be linked to greater transmissibility, lower vaccine efficiency, and an increased risk of reinfection. Globally, the number of nations reporting SARS-CoV-2 Omicron variant of concern (VOC) infections continues to rise, with a total of 352 confirmed cases reported by 27 countries as of December 1, 2021. It is uncertain if the Omicron COVID variation is more transmissible or severe than the Delta variant form. The purpose of this study was to compare the binding affinity of SARS-CoV-2 delta and omicron variants with ACE2 by using a variety of computational tools to compare the binding affinity of Wuhan-Hu-1 with delta and omicron variants.

## Methodology

### Data retrieval

The FASTA sequence of the spike protein of SARS-CoV-2 of Wuhan-Hu-1 (wild type) was obtained from Uniport ^13^. (Accession no: P0DTC2). The delta variant spike protein (accession no. QWK65230.1) was obtained from ViPR (Virus Pathogen Resource) ^14^. The omicron complete genome (R40B60 BHP 3321001247/2021) was obtained from GSAID ^15^ and the genome sequence was translated to protein sequence using the expasy translate programme ^16^. The translated sequence was used to select the Omicron spike protein.

### Analysis of physicochemical parameter

The Wuhan-Hu-1 (wild type), delta, and omicron variant sequences were analysed using the ExPASy ProtParam online tool. ProtParam calculates the molecular weight, theoretical pI, amino acid composition, atomic composition, extinction coefficient, anticipated half-life, instability index, aliphatic index, and grand average of hydropathicity (GRAVY).

### Prediction of secondary structural changes in spike protein

GOR IV ^17^ was used to predict the secondary structure of the Wuhan-Hu-1, delta, and omicron variants. The GOR (Garnier–Osguthorpe–Robson) tool uses information theory and Bayesian statistics to analyse secondary protein structure. The goal of combining multiple sequence alignments using GOR is to gain knowledge for improved secondary structure differentiation.

### Identification of Conserved Residues and Mutation

Clustal Omega ^18^ a bioinformatics programme, was used to align the Wuhan-Hu-1 (wild type) sequence with variants of delta and omicron sequences. The box shade application was used to create the alignment figure.

### Intrinsically unstructured protein prediction

Intrinsic disorder regions (IDRs) are locations in physiological contexts that have a dynamic ensemble of conformations that do not acquire a stable three-dimensional structure. The Wuhan-Hu-1, Delta variant, and Omicron variant sequences were predicted using the Predictor of Natural Disordered Regions (PONDR) ^19^.

### Prediction of Protein Stability

Wuhan-Hu-1, Delta, and Omicron sequences were predicted using I-Mutant3.0 ^20^. It is a support vector machine (SVM)-based tool for predicting protein stability changes resulting from single point mutations. It may be used to predict the sign of the protein stability change caused by mutation, as well as as a regression estimator to predict the associated G values. Protein structure dynamics and flexibility are also important aspects of protein function. PredyFlexy ^21^ was used to predict extremely flexible protein structures to better understand the features of their protein.

### SIFT for prediction of the effect of nsSNPs on protein function

The Wild type, Delta, and Omicron variants are checked whether mutation impacts protein function through SIFT tool ^22^. SIFT predicts whether an amino acid substitution affects protein function based on sequence homology and the physical properties of amino acids.

### Prediction of disease-associated

VarSite ^23^ is a web server mapping known disease□associated variants from UniProt and ClinVar, together with natural variants from gnomAD, onto protein 3D structures in the Protein Data Bank. The spike protein variants undergo mutation by interaction with human ACE2 protein. The mutation changes of SARS-CoV-2 were predicted using varsite.

### Mutagenesis analysis

The PDB file contains the crystal structure of the SARS-CoV-2 spike receptor-binding domain coupled to ACE2 (6M0J) ^24^. The complex is composed of two protein chains: SARS-CoV-2 RBD (Chain B) and human ACE2 (Chain A). Chain B’s SARS-CoV-2 RBD was separated and utilised for further mutagenesis study. The SARS-CoV-2 RBD mutations were introduced using the Pymol mutagenesis wizard programme at the appropriate delta and omicron mutated positions for each residue and the whole RBD.

### Protein-Protein docking of mutated RBD and human ACE2

After preparing the hACE2 and SARS-CoV-2 spike protein receptors using both full spike protein and RBD, all potentially docked molecules were analysed using the HEX docking programme ^25^. Docking parameters were set as follows: Type of correlation: Only the shape dimension is 0.6, the receptor range is 180, the ligand range is 180, the distance range is 40, and the box size is ten. OPLS minimisation as a post-processing step. Following that, the best docking results were achieved using the HEX programme.

## Results and Discussion

The current global pandemic coronavirus infection (COVID-19), which began in late December 2019 in Wuhan, China, is suspected to be caused by the SARS-CoV-2 coronavirus. The Centers for Disease Control and Prevention has categorised SARS-CoV-2 variants as variants of interest, variants of concern, and variants of high importance (CDC).

Several SARS-CoV-2 variants have been identified, posing a long-term infection risk in immunocompromised individuals ^26^. The “variant of concern” (VOC) SARS-CoV-2 (which generates COVID-19) refers to viral variants in which mutations in the spike protein receptor-binding domain (RBD) drastically increase binding affinity in the RBD-hACE2 complex while also being connected to rapid transmission in human populations ^27^. A variety of computational approaches are utilised in this study to compare the currently categorised omicron variant to the delta variant to get insight into its characteristics and binding affinity for the ACE2 protein.

### Multiple alignment of Delta and Omicron variant with Wuhan-Hu-1

The omicron variation includes 30 mutations in the Spike protein, half of which are in the receptor-binding domain, according to the multiple alignments **(Figure 1)**. From a previous study, it is observed that RBD T470-T478 loop and Y505 as viral determinants for specific recognition of SARS-CoV-2 RBD by ACE2 ^28^. T478 is a common mutation seen in Delta and Omicron variants **(Figure 2)**. RBD has the potential to be developed into an efficient and safe subunit vaccine against SARS-CoV-2 due to its ability to produce very robust nAb responses. Numerous mutations in the spike protein’s receptor-binding region in Omicron compared to the Delta variant **(Figure 3)** suggests that the Omicron variant may be immunologically resistant to antibody-mediated protection **(Table 1)**.

**Table 1:** Spike protein mutation in Delta and Omicron variant compared to wild type (Wuhan-Hu-1)

| Variant | Sequence ID | Mutation |
| --- | --- | --- |
| Wuhan-Hu-1<br>(wild type) | NCBI ID:P0DTC2 | - |
| Delta Variant<br>(B.1.617.2) | NCBI: QWK65230.1 | T19R, G142D, Δ156-157, R158G, Δ213-214, **L452R**, **T478K**, D614G, P681R, D950N |
| Omicron<br>(B.1.1.529) | GSAID ID:<br>R40B60_BHP_3321001247/2021 | A67V, Δ69-70, T95I, G142D, Δ143-145, N211I, L212V, ins213-214RE, V215P, R216E, **G339D**, **S371L**, **S373P**, **S375F**, **K417N**, **N440K**, **G446S**, **S477N**, **T478K**, **E484A**, **Q493R**, **G496S**, **Q498R**, **N501Y**, **Y505H**, T547K, D614G, H655Y, N679K, P681H, N764K, D796Y, N856K, Q954H, N969K, L981F |

\* Receptor binding domain (**residues 319-541**) are marked as bold in both Delta and Omicron variant. Δ represents deletion, ins represents insertion

**Figure 1:**
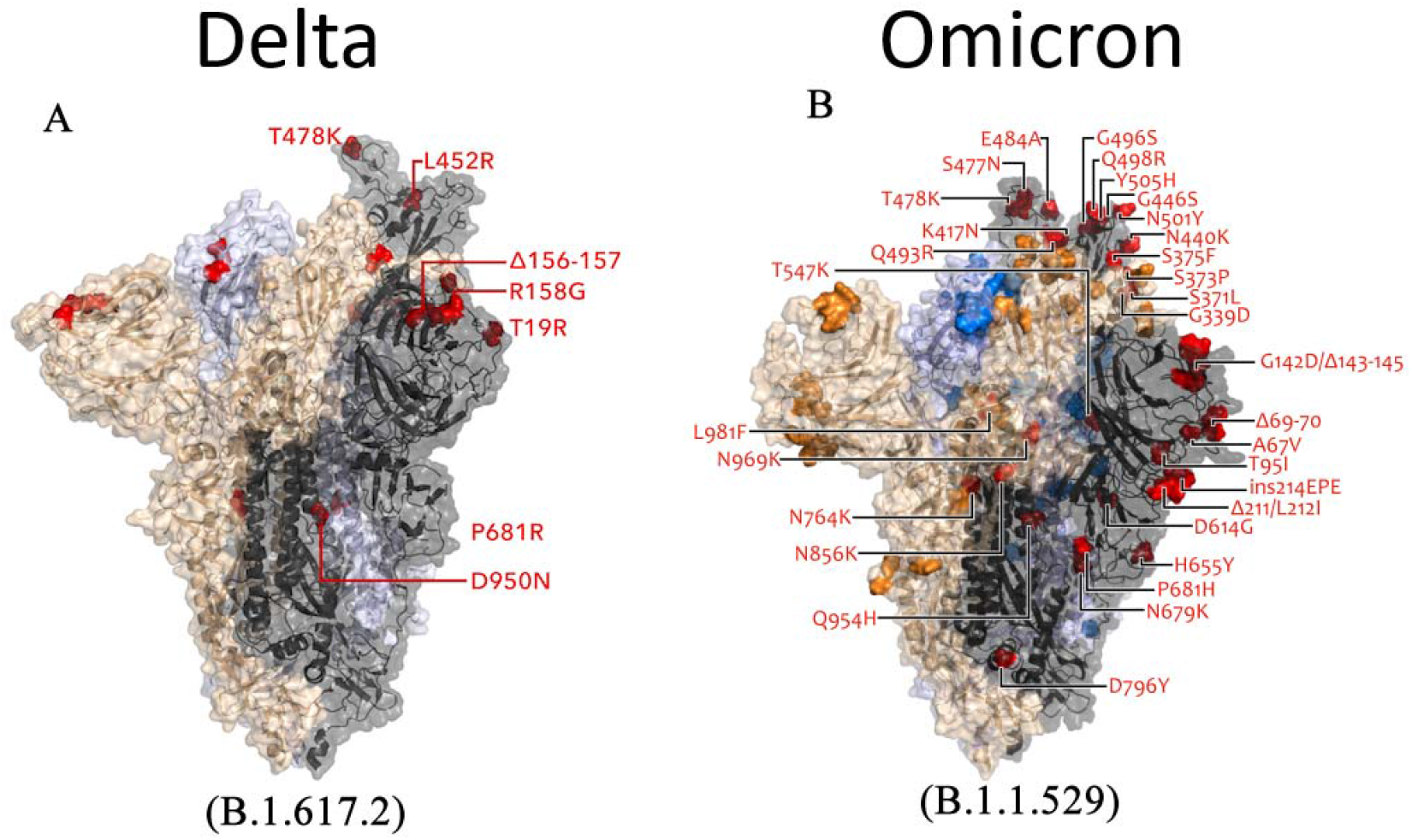
A comparison of Delta and Omicron variant spike mutation (Image source: Modified from COVID-19 Genomics UK Consortium).

**Figure 2:**
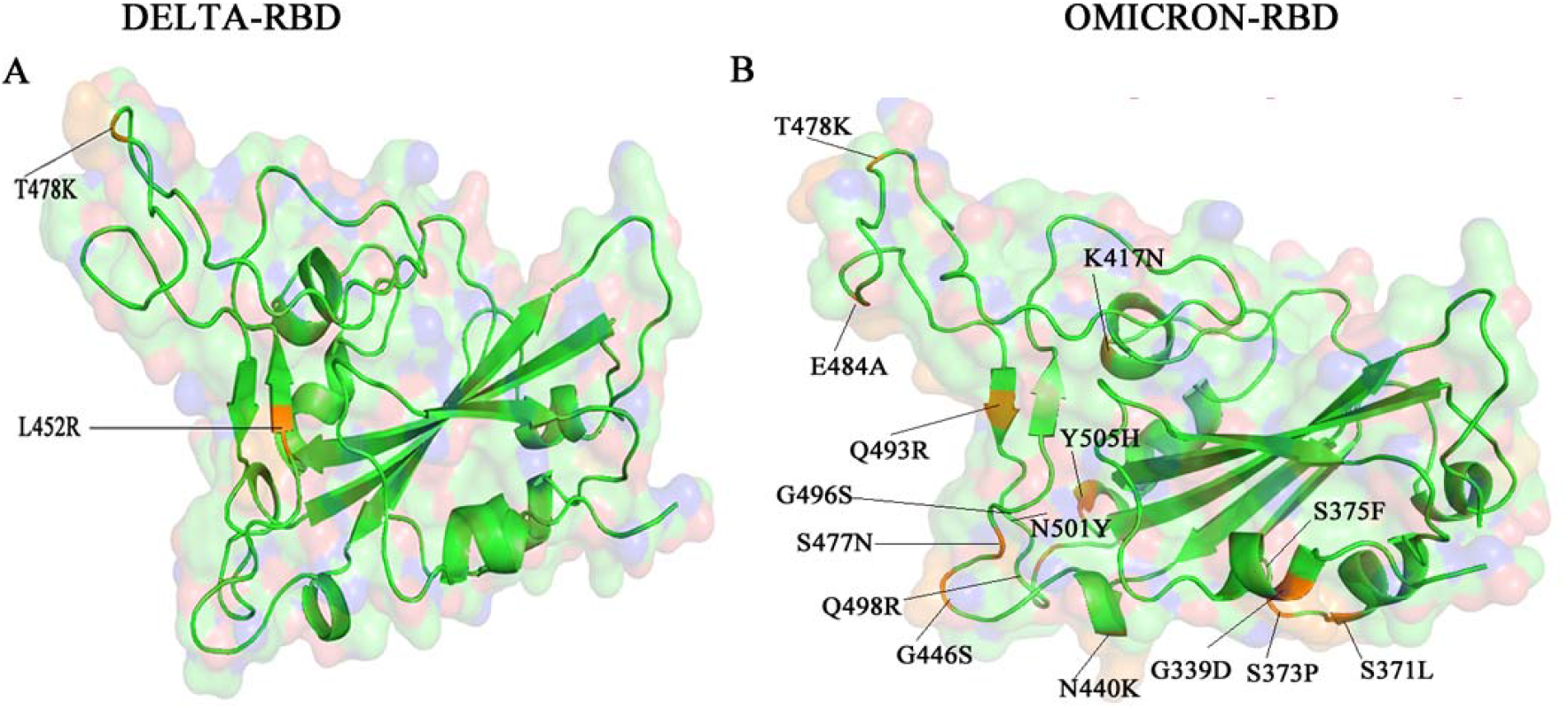
A comparison of Delta and Omicron variant mutation in Receptor Binding Domain (RBD). The mutation is marked in yellow colour. Delta-RBD has only two mutations whereas Omicron-RBD has 15 mutations.

**Figure 3:**
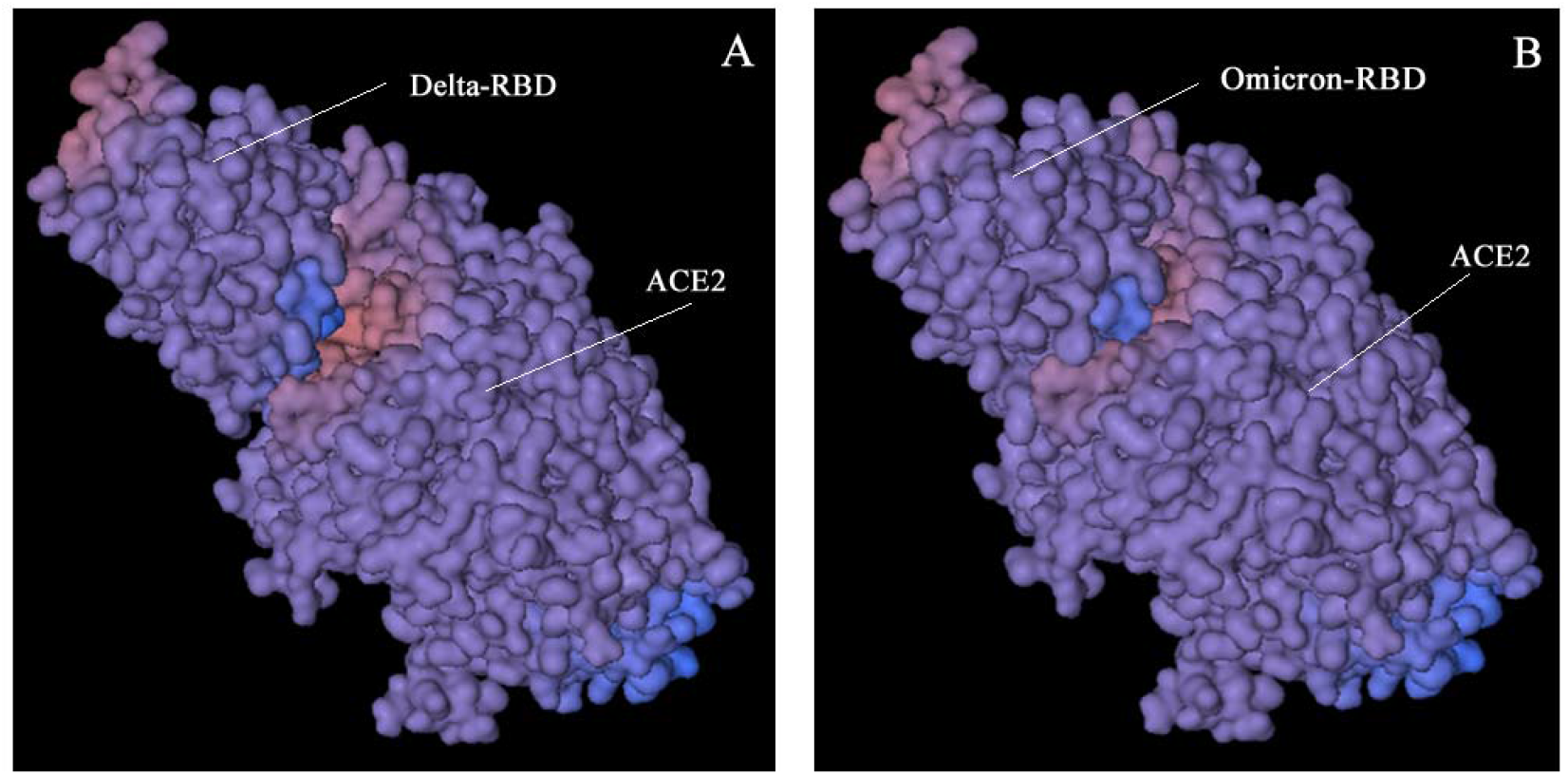
Docking between **(A)** Delta-RBD and **(B)**Omicron-RBD with ACE2.

### Determination of physical parameters of the proteins

While Wuhan-Hu-1 has 1273 amino acids, the delta variant has 1271 and the omicron variant has 1270; nevertheless, due to sequence loss, both the delta and omicron variants have a few fewer residues than the wild type. A protein’s isoelectric point (pI) is the pH value at which its surface is completely charged but its net charge is zero. A pI value of more than 7 indicates that the protein is alkaline, whereas a value less than 7 indicates that it is acidic. The molecular weight of Wuhan-Hu-1 is 141178.47 with a theoretical pI of 6.24, the delta variant is 140986.31 with a theoretical pI of 6.78, and the omicron variant is 141328.11 with a theoretical pI of 7.14. Despite having three fewer amino acids than Wuhan-Hu-1, the omicron variant has a higher molecular weight and theoretical PI than the delta variant and Wuhan-Hu-1. In the current investigations, the omicron variant is expected to have an alkaline pI, while the delta and Wuhan-Hu-1 variants are expected to have an acidic pI. According to previous research ^29^ a stability score of less than 40 indicates that the protein structure is stable. A value of 40 or above suggests that the protein is structurally unstable. In our research, the range remained 32.81-34.69, indicating the great stability of all SARS-CoV-2 spike proteins. The average extinction coefficient is 11238.61, which indicates how much light the protein can absorb at 280 nm. The aliphatic index measures the volume of a protein that is filled by aliphatic amino acids on the side chain, such as alanine. A high aliphatic index of 84.50 to 84.95 indicates that the protein is temperature stable across a wide temperature range. The greater a protein’s aliphatic index, the more thermostable it is. The degree to which amino acids in a protein sequence are hydrophobic or hydrophilic is referred to as hydropathicity. A protein with a low GRAVY (Grand average of hydrophobicity) value is nonpolar and has a stronger affinity for water, indicating that it is intrinsically hydrophilic.

Primary structural study indicates a set of features shared by all SARS-CoV-2 variants. According to the amino acid composition of the omicron variant, there is an increase in the following amino acid compositions compared to the delta variant: Arginine (Arg), Lysine (Lys), Aspartic acid (Asp), and Glutamic acid (Glu), indicating that the omicron has more charged residues that contribute to salt bridge formation and that charged residues are exposed to a much greater degree.

The higher amino acid composition of Phenylalanine (F), Isoleucine (I) in the omicron spike protein, when compared to the delta variant, suggests that the omicron spike protein includes more hydrophobic amino acids, which may be due to its positioning inside the protein core. When compared to the delta version, the omicron variant’s amino acid composition is low in polar amino acids such as Asparagine (N), Glutamine (Q). Omicron RBD is high in non-polar amino acids such as Leucine (L), Phenylalanine (F), and Proline (P) ^30^. These residues are located inside the protein core and are thus inaccessible to the solvent **(Table 2)**.

**Table 2:** Amino acid composition comparison between Delta and Omicron variant with reference to wild type (wuhan-Hu-1)

|  | Wuhan-Hu-1<br>-Whole Spike | Wuhan-Hu-1<br>RBD | Delta-Whole<br>Spike | Delta-RBD | Omicron-Whole<br>Spike | Omicron-RBD |
| --- | --- | --- | --- | --- | --- | --- |
| **Ala (A)%** | 6.2 | 5.2% | 6.2 | 5.2% | 6.2 | 5.7% |
| **Arg (R)%** | 3.3 | 4.8 | 3.5 | 5.2% | 3.5 | 5.7% |
| **Asn (N)%** | 6.9 | 9.2% | 7.0 | 10.0% | 6.5 | 10.0% |
| **Asp (D)%** | 4.9 | 3.9% | 4.8 | 3.9% | 4.9 | 4.4% |
| **Cys (C)%** | 3.1 | 3.9% | 3.1 | 3.9% | 3.1 | 3.9% |
| **Gln (Q)%** | 4.9 | 3.1% | 4.9 | 3.1% | 4.6 | 2.2% |
| **Glu (E)%** | 3.8 | 3.1% | 3.7 | 3.1% | 3.9 | 2.6% |
| **Gly (G)%** | 6.4 | 6.6% | 6.5 | 7.0% | 6.2 | 5.7% |
| **His (H)%** | 1.3 | 3.1% | 1.3 | 0.4% | 1.4 | 0.9% |
| **Ile (I)%** | 6.0 | 3.9% | 6.0 | 3.9% | 6.1 | 3.9% |
| **Leu (L)%** | 8.5 | 6.1% | 8.4 | 6.1% | 8.4 | 7.0% |
| **Lys (K)%** | 4.8 | 5.2% | 4.9 | 5.7% | 5.3 | 6.1% |
| **Met (M)%** | 1.1 | 0.0% | 1.1 | 0.0% | 1.1 | 0.0% |
| **Phe (F)%** | 6.0 | 7.0% | 6.0 | 7.4% | 6.2 | 7.9% |
| **Pro (P)%** | 4.6 | 5.7% | 4.5 | 5.7% | 4.6 | 6.1% |
| **Ser (S)%** | 7.8 | 7.4% | 7.8 | 7.4% | 7.6 | 6.6% |
| **Thr (T)%** | 7.6 | 5.7% | 7.5 | 5.7% | 7.4 | 5.2% |
| **Trp (W)%** | 0.9 | 0.9% | 0.9 | 0.9% | 0.9 | 0.9% |
| **Tyr (Y)%** | 4.2 | 6.6% | 4.2 | 6.6% | 4.3 | 6.6% |
| **Val (V)%** | 7.6 | 8.7% | 7.6 | 8.7% | 7.6 | 8.7% |
| **Pyl (O)%** | 0.0 | .0 | 0.0 | 0.0% | 0.0 | 0.0 |
| **Sec (U)%** | 0.0 | 0.0 | 0.0 | 0.0% | 0.0 | 0.0 |

### Prediction of secondary structural changes

Omicron has a higher fraction of alpha-helix structure (23.46%) than delta variant (22.03%), but less extended strand and random coil structure **(Table 3)**. Because it is largely made up of alpha-helices, the evidence suggests that the omicron variant protein is structurally highly stable. The Omicron form of RBD has a greater alpha helix composition than the Delta variation in secondary structure prediction, suggesting that the RBD has a stable structure, however, the random coil composition is slightly increased in Omicron ^31^.

**Table 3:**
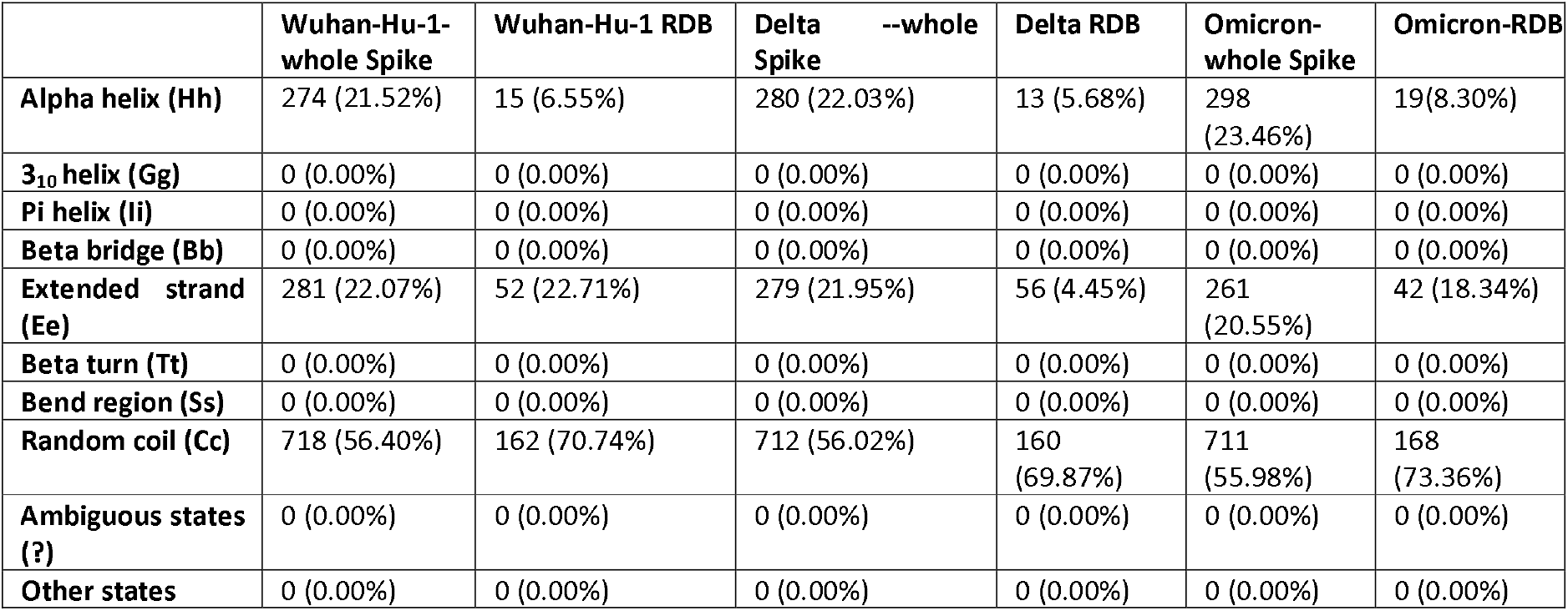
Secondary structure prediction and comparison of Delta and Omicron variant with reference to wild type (Wuhan-Hu-1)

### Intrinsically disordered Prediction

Disordered areas of viral proteins are linked to viral pathogenicity and infectivity. PONDR® VLXT was used to predict the intrinsic disorder of Wuhan-Hu-1, Delta, and Omicron variants. Residues with anticipated disorder scores more than 0.5 are regarded inherently disordered, while residues with expected disorder values between 0.2 and 0.5 are considered flexible. According to the prediction, the Omicron variant has a less disordered area than the Delta variant and the wild type. We observed that disordered regions in entire spike protein as well as RBD in Omicron exhibit disorder-to-order transition when compared to Delta variant and wild type. According to prior research from the cryo-EM structure of Wuhan-Hu-1, the T470-F490 loop and Q498-Y505 within RBD are key contacting elements that interact with RBD and ACE2 ^28^. The disorder prediction ranges from 468-473 with residues ISTEIYQA in Wuhan-Hu-1-RBD, 469-471 with residues EIY in Delta variant-RBD, and there are no disorder residues predicted in this region in Omicron Variant-RBD. This implies that there is a chance of disorder-order transition between region 468-473 of spike protein, which could be important in the influence of disordered residues/regions on spike protein stability and binding to ACE2. **(Table 4)**.

**Table 4:**
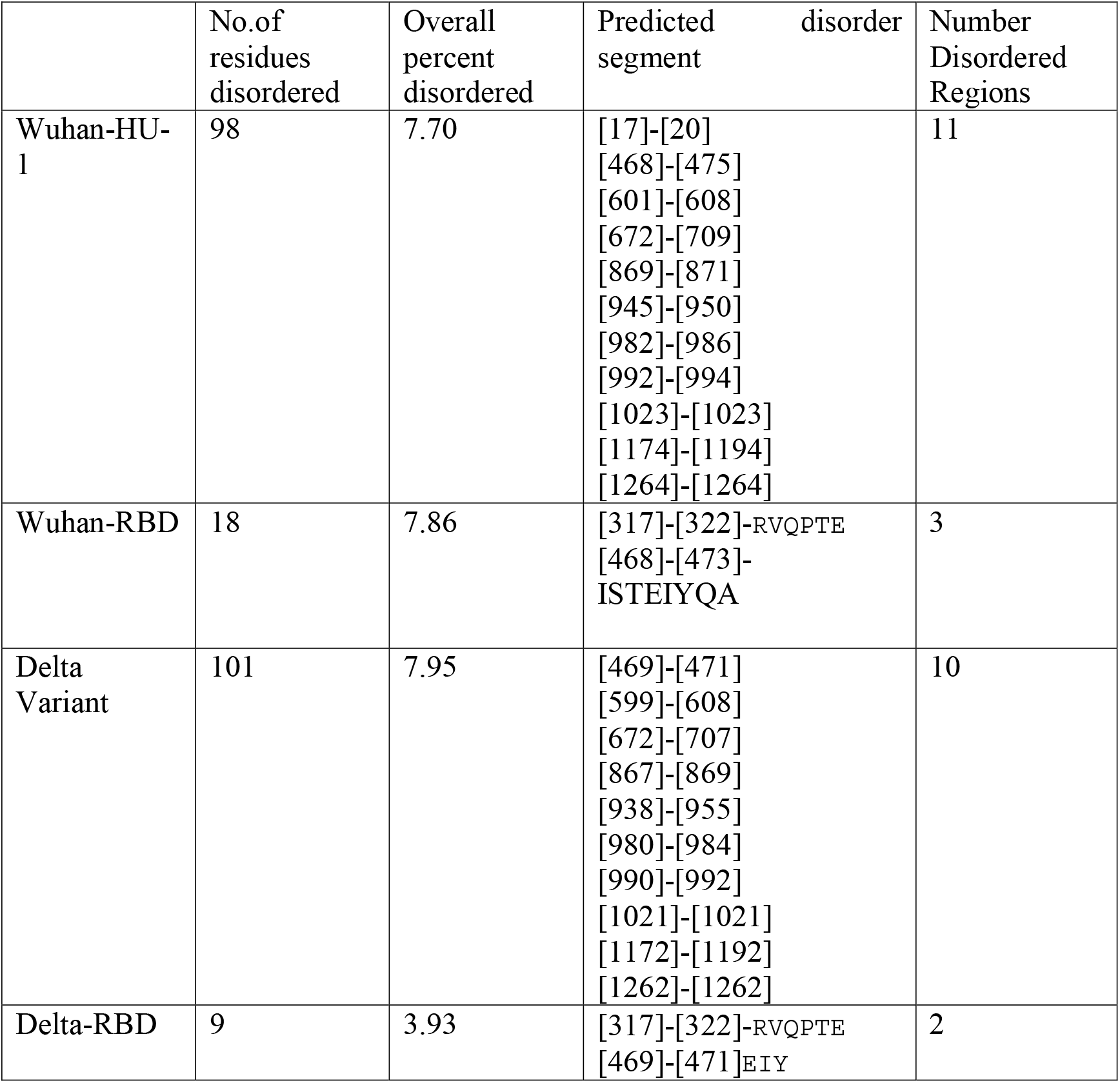
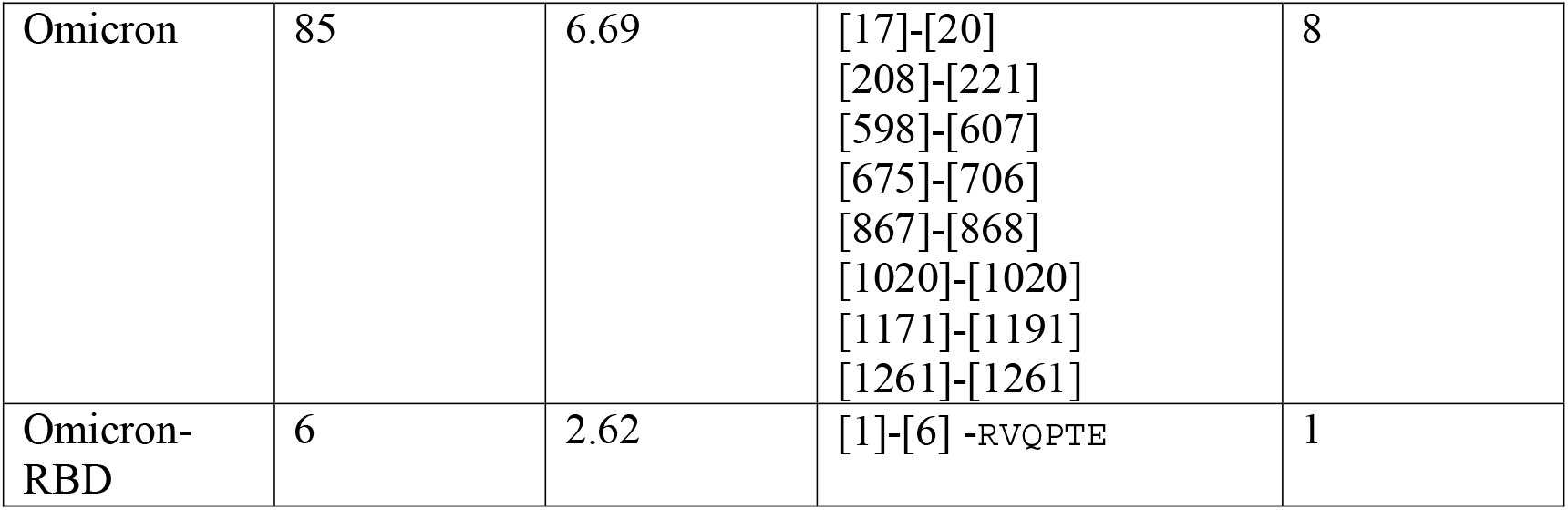
Intrinsically disordered prediction

### Prediction of Protein stability changes upon mutation

An I-Mutant protein stability study predicted that all amino acid modifications in the delta variant reduce spike protein stability. Except for the N501Y mutation ^32^, which is expected to improve the stability of the spike protein, all amino acid changes in the omicron variant result in a decrease in stability **(Table 5)**. SIFT analysis revealed that, whereas the delta variant D950N impairs protein function, other mutations are tolerated. The N211I, Y505H, and N764K mutations in the omicron variant impair protein function, although other variants are tolerated **(Table 5)**. Although the RBD L452R and T478 delta mutations are tolerated, they reduce protein stability and increase disease risk. There are 15 Omicron variant mutations in RBD, the N501Y mutation being one of them. It is tolerated and enhances protein stability; however, it is disease-prone. Other mutations that decrease protein stability and increase disease risks, such as G339D, S371L, S373P, S375F, N440K, G446S, T478K, G496S, and Q498R, are tolerated. Tolerable mutations include K417N, S477N, E484A, Q493R, and Q498R, which decrease protein stability and increase disease vulnerability. Protein function is impaired by Y505H mutations, resulting in decreased protein stability and an increased risk of disease **(Table 6)**.

**Table 5:** Protein Stability analysis for single point mutation using I-Mutant and SIFT tools. The disease propensity predicted using Varsite

| Variant | AA change | I-Mutant | SIFT | Varsite -disease propensity (>1 more disease prone) |
| --- | --- | --- | --- | --- |
| **Delta (B.1.617.2)** |  |  |  |  |
|  | T19R | Decrease stability<br>-0.40 | TOLERATED (0.10) | 1.16 |
|  | G142D | Decrease stability<br>-1.15 | TOLERATED (0.18) | 1.93 |
|  | R158G | Decrease stability<br>-1.65 | TOLERATED (0.08) | 0.99 |
|  | **L452R** | Decrease stability<br>-1.85 | TOLERATED (0.38) | 2.36 |
|  | **T478K** | Decrease stability<br>-0.74 | TOLERATED (0.75) | 1.04 |
|  | D614G | Decrease stability<br>-0.93 | TOLERATED (0.62) | 1.00 |
|  | P681R | Decrease stability<br>-0.90 | TOLERATED (0.33) | 0.89 |
|  | D950N | Decrease stability<br>-0.43 | AFFECT PROTEIN FUNCTION (0.00) | 0.99 |
| **Omicron** |  |  |  |  |
|  | A67V | Decrease stability<br>-0.01 | TOLERATED (0.58) | 0.75 |
|  | T95I | Decrease stability<br>-0.78 | TOLERATED (0.14) | 0.66 |
|  | G142D | Decrease stability<br>-1.15 | TOLERATED (0.18) | 1.93 |
|  | N211I | Increase stability<br>1.06 | AFFECT PROTEIN FUNCTION (0.00) | 1.34 |
|  | L212V | Decrease stability<br>-1.75 | TOLERATED (0.33) | 0.47 |
|  | V213P | Decrease stability<br>-1.69 | TOLERATED (0.26) | V213P, is not shown on the histogram as in involves 2 base changes in the corresponding DNA codon |
|  | R214E | Decrease stability<br>-0.71 | TOLERATED (0.08) | R214E, is not shown on the histogram as in involves 2 base |

|  |  |  |  | changes in the corresponding DNA codon. |
| --- | --- | --- | --- | --- |
|  | **G339D** | Decrease stability<br>-1.16 | TOLERATED (0.88) | 1.93 |
|  | S371L | Decrease stability<br>-0.22 | TOLERATED (1.00) | 1.08 |
|  | S373P | Decrease stability<br>-0.53 | TOLERATED (0.38) | 1.32 |
|  | S375F | Decrease stability<br>-0.33 | TOLERATED (0.07) | 1.00 |
|  | K417N | Decrease stability<br>-0.42 | TOLERATED (0.56) | 0.76 |
|  | N440K | Decrease stability<br>-0.50 | TOLERATED (0.70) | 1.04 |
|  | G446S | Decrease stability<br>-1.49 | TOLERATED (0.79) | 1.34 |
|  | S477N | Decrease stability<br>-0.45 | TOLERATED (0.84) | 0.46 |
|  | T478K | Decrease stability<br>-0.74 | TOLERATED (0.75) | 1.04 |
|  | E484A | Decrease stability<br>-0.79 | TOLERATED (0.51) | 0.66 |
|  | Q493R | Decrease stability<br>-0.17 | TOLERATED (0.51) | 0.65 |
|  | G496S | Decrease stability<br>-1.22 | TOLERATED (0.22) | 1.34 |
|  | Q498R | Decrease stability<br>-0.17 | TOLERATED (0.39) | 0.65 |
|  | N501Y | Increase stability<br>0.15 | TOLERATED (0.09) | 1.22 |
|  | **Y505H** | Decrease stability<br>-1.49 | AFFECT PROTEIN<br>FUNCTION (0.03) | 1.11 |
|  | T547K | Decrease stability<br>-1.05 | TOLERATED (0.83) | 1.04 |
|  | D614G | Decrease stability<br>-0.93 | TOLERATED (0.62) | 1.00 |
|  | H655Y | Increase stability<br>0.08 | TOLERATED (0.50) | 0.76 |
|  | N679K | Decrease stability<br>-0.32 | TOLERATED (0.53) | 1.04 |
|  | P681H | Decrease stability<br>-1.27 | TOLERATED (0.17) | 0.94 |
|  | N764K | Decrease stability<br>-0.21 | AFFECT PROTEIN<br>FUNCTION (0.00) | 1.04 |
|  | D796Y | Decrease stability<br>-0.09 | TOLERATED (1.00) | 1.44 |
|  | N856K | Decrease stability<br>-0.38 | TOLERATED (0.08) | 1.04 |
|  | Q954H | Decrease stability<br>-0.86 | TOLERATED (0.08) | 0.49 |
|  | N969K | Decrease stability<br>-0.63 | TOLERATED (0.09) | 1.04 |
|  | L981F | Decrease stability<br>-1.24 | TOLERATED (0.27) | 0.64 |

**Table 6:**
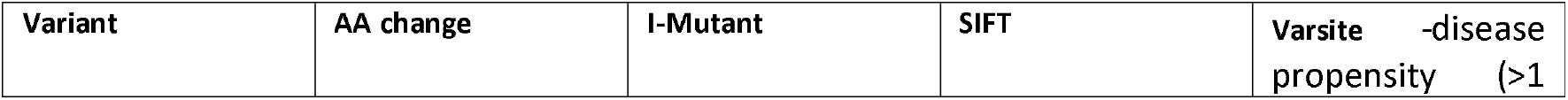
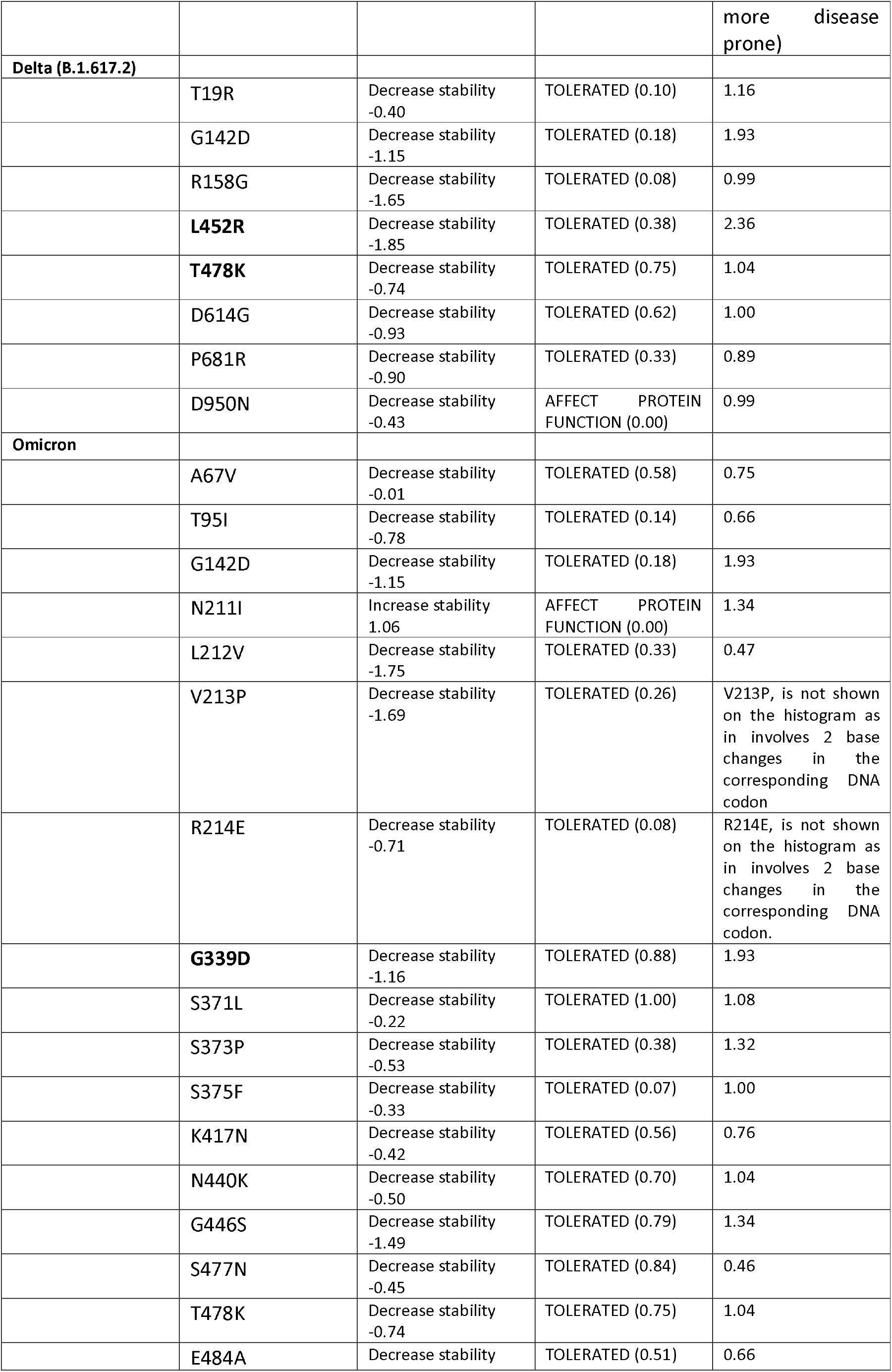
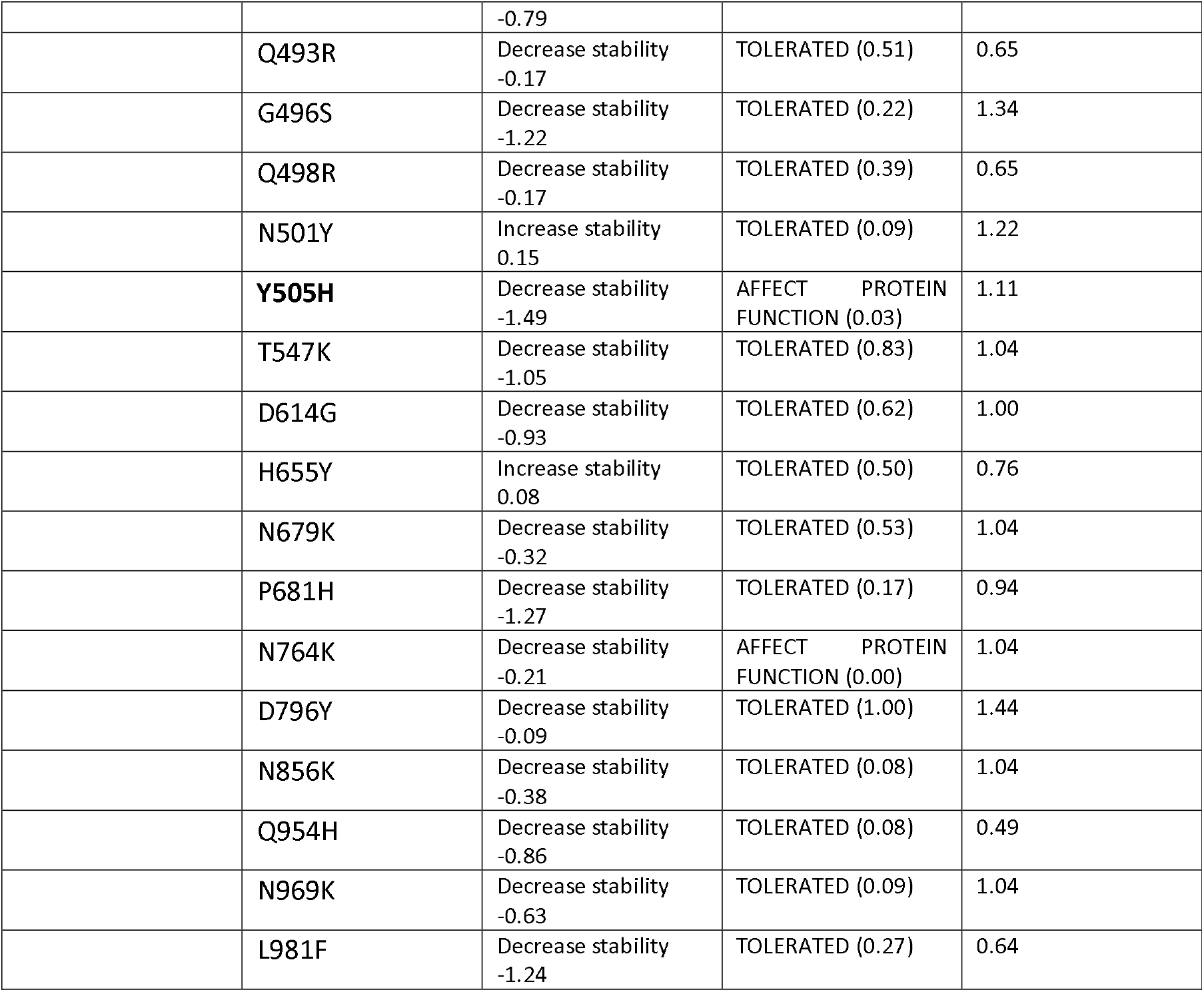
Protein stability analysis using PremPs

The large type I transmembrane S glycoprotein on the viral envelope and the homologous receptor on the surface of host cells enable membrane fusion. The S glycoprotein’s exposed surface not only allows membrane fusion but also drives host immune responses, making it a great target for neutralising antibodies ^33^. Cleavage at the S1/S2 site results in the formation of a surface subunit S1, which attaches the virus to the host cell surface receptor, and a transmembrane component S2, which allows the viral and host cell membranes to merge. The S2 subunit of the transmembrane is made up of an N-terminal hydrophobic fusion peptide (FP), two heptad repeats (HR1 and HR2), a transmembrane domain (TM), and a cytoplasmic tail (CT), in the following order: FP-HR1-HR2-TM-CT ^34^. The amino acid residues Y695, I923, S982, V1189, F1220, and I1221 in Wuhan-Hu-1 are extremely flexible. Residues I921, S980, V1187, F1218, and I1219 are extremely variable in the Delta variant. The I920, S979, V1186, F1217, and I1218 residues are particularly flexible in the Omicron variant, as predicted by PredyFlexy. Flexible prediction and local structure prediction from sequence show that the heptapeptide repeat sequence 1 (HR1) (912–984 residues), HR2 (1163–1213 residues), and TM domain (1213–1237 residues) of the S2 subunit are very flexible in both the Delta and Omicron variants.

### SARS-CoV-2 RBD-hACE2 docking

Understanding the SARS-CoV-2 virus’s receptor recognition mechanism is critical since it governs the virus’s infectivity, host range, and pathogenesis. The binding affinity of SARS-CoV-2 variants of RBD to ACE2 differs because of minor variations in ACE2 interactions. In this study, the PDB (6M0J) crystal structure of the SARS-CoV-2 spike receptor-binding domain associated with ACE2 was employed. The RBD of SARS-CoV-2 was isolated from ACE2 and used for protein-protein docking. Hex was used to dock SARS-CoV-2 RBD and ACE2, and its docking score (−500.37) is for Wuhan-Hu-1 (Wild type), which was utilised to compare the docking energies of Delta and Omicron. The pymol mutagenesis wizard was used to add mutations into the Delta and Omicron versions. Docking was performed using hACE2 between the Delta and Omicron Variant **(Figure 4)**. The docking score for the Omicron variation is the highest (−539.81), while the Delta variant is the lowest (−529.62). This suggests that the Omicron variant is more responsive to hACE2 than the Delta variant, indicating a higher potential for transmission **(Table 7)**. In addition, the impact of each changed residue on hACE2 affinity was investigated. The highest binding affinity score of all 15 RBD mutations is Q493R (−581.53), followed by N501Y (−560.81), S371L (−54.34), S373P (−541.87), S375F (−530.07), Q498R (−527.38), and T478 (−517.03) **(Table 8)**. For the Delta version, only two mutations were found in RBD, with L452R (−517.52) having the highest binding affinity, followed by T478 (−517.03). Point mutations at key residues have a significant impact on the interaction with ACE2. SARS-CoV-2 interacts with hACE2 through its C-terminal domain (SARS-CoV-2-CTD), indicating that it has a higher affinity for the receptor. In SARS-CoV-2-CTD, E484 forms ionic contacts with K31, increasing receptor affinity. Previous research has found that the single mutation E484 in the viral spike (S) protein (which is shared by the Beta and Gamma VOCs as well as the Mu VOI) may be critical in avoiding vaccination immunity; variants with the E484 mutation have demonstrated resistance to neutralising antibodies generated by prior infection ^35^. The E484A mutation (478.49) has binding affinity in the Omicron variant may result in enhanced hACE2 binding. The Omicron form of ACE2 binds more strongly to SARS-CoV-2 than the Delta variant of hACE2.

**Table 7:** Docking analysis of spike protein with ACE2 using HEX software

|  | Docking Energy |
| --- | --- |
| Wildtype (wuhan-Hu-1)-Ace2 | -500.37 |
| omicron | -539.81 |
| delta | -529.62 |

**Table 8:** Docking analysis of Wuhan-RBD, Delta-RBD, Omicron-RBD with ACE2 using HEX software

| Variant with ACE2 | RBD Mutation | Docking energy |
| --- | --- | --- |
| Wild |  | -500.37 |
| Delta | L452R | -517.52 |
|  | T478K | -517.03 |
| Omicron |  |  |
|  | G339D | -507.06 |
|  | S371L | -549.34 |
|  | S373P | -541.87 |
|  | S375F | -530.07 |
|  | K417N | -500.42 |
|  | N440K | -496.38 |
|  | G446S | -503.18 |
|  | S477N | -500.05 |
|  | T478K | -517.03 |
|  | E484A | -478.49 |
|  | Q493R | -581.53 |
|  | G496S | -505.58 |
|  | Q498R | -527.38 |
|  | N501Y | -560.81 |
|  | Y505H | -502.24 |

The existence of a high number of Omicron variant mutations is also a hallmark of the variants, indicating that viral evolution in immunocompromised persons may have played a significant role in their development. Because many people worldwide suffer from inherent or induced immunosuppression, the relationship between immunosuppression and the generation of highly transmissible or pathogenic SARS-CoV-2 variants must be investigated further and mitigation strategies devised.

## Conclusion

Both the Omicron and Delta variants were investigated in this study using several computational tools and a computational saturation mutagenesis model, examining structural, sequence-driven, and dynamic changes that effect overall protein stability were examined. According to the findings of this study, large changes in the RBD region of the Omicron variant contribute to higher binding with hACE2, which may result in a higher transmission rate when compared to the Delta variant.

## Acknowledgments

We acknowledge Faculty of Health and Life Sciences, Management and Science University, Malaysia; Division of Cardiovascular Medicine, Radcliffe Department of Medicine, Wellcome Centre for Human Genetics, University of Oxford, Oxford, UK and Department of Physiology, Anatomy and Genetics, University of Oxford, Oxford, UK. We also acknowledge GISAID (https://www.gisaid.org/) for facilitating open data sharing.

